# A Nostalgia Brain-Music Interface for Enhancing Nostalgia, Well-Being, and Memory Vividness in Young and Elderly Individuals

**DOI:** 10.1101/2024.10.29.620793

**Authors:** Yuna Sakakibara, Tomohiro Kusutomi, Sotaro Kondoh, Takahide Etani, Saori Shimada, Yasuhiko Imamura, Yasushi Naruse, Shinya Fujii, Takuya Ibaraki

## Abstract

Music-evoked nostalgia has the potential to assist in recalling autobiographical memories and enhancing well-being. However, nostalgic music preferences vary from person to person, presenting challenges for applying nostalgia-based music interventions in clinical settings, such as a non-pharmacological approach. To address these individual differences, we developed the Nostalgia Brain-Music Interface (N-BMI), a neurofeedback system that recommends nostalgic songs tailored to each individual. This system is based on prediction models of nostalgic feelings, developed by integrating subjective nostalgia ratings, acoustic features and in-ear electroencephalographic (EEG) data during song listening. To test the effects of N-BMI on nostalgic feelings, well-being, and memory recall, seventeen elderly and seventeen young participants took part in the study. The N-BMI was personalized for each individual, and songs were recommended under two conditions: the “nostalgia condition”, where songs were selected to enhance nostalgic feelings, and the “control condition”, to reduce nostalgic feelings. We found nostalgic feelings, well-being, and memory vividness were significantly higher after listening to the recommended songs in the nostalgia condition compared to the control condition in both groups. This indicates that the N-BMI enhanced nostalgic feelings, well-being, and memory recall across both groups. The N-BMI paves the way for innovative therapeutic interventions, including non-pharmacological approaches.

## Introduction

As the global population continues to grow and age, the incidence of dementia—a clinical syndrome characterized by progressive cognitive decline—has become an increasingly pressing issue. The risk of developing dementia increases with age, leading to a projected rise in the number of individuals affected^1^. According to the Global Burden of Disease Study 2019, this increase is expected to result in approximately 151 million people living with dementia by 2050^2^. Despite the growing prevalence of dementia, pharmacological treatments remain challenging due to the disease’s complex nature and associated side effects.

As a result, non-pharmacological approaches have gained attention as low-risk, accessible options for dementia care and rehabilitation^3,4^. Among these, music therapy may be effective in two perspectives: memory retention and psychological well-being^5^. One notable aspect of music’s impact on dementia care is its potential to aid in memory recovery. Memory loss is a primary symptom of dementia, as well as mild cognitive impairment (MCI), a condition often seen as a precursor to dementia^6,7^. Music has been shown to have a unique ability to preserve long-term memory, even in individuals with Alzheimer’s disease, where memory can remain intact even as other cognitive functions deteriorate^8,9^. Music is particularly powerful in evoking autobiographical memories^10^, a phenomenon observed even in dementia patients with significant memory loss. Studies have shown that listening to music can trigger related memories in patients with amnesia and dementia^11–16^. These findings suggest that music could play a crucial role in slowing the progression of memory loss in dementia patients^17^.

Additionally, music has the potential to enhance the emotional well-being of dementia patients. Behavioral and psychological symptoms of dementia (BPSD), such as depression and anxiety, are significant concerns in dementia care^18^. Music therapy has been shown to alleviate these symptoms, improving psychological health and overall well-being^19–21^.

In particular, nostalgic music may be especially effective in preserving memory and enhancing psychological health in dementia patients. Nostalgia is a bittersweet emotional experience associated with recollections of the past, serving as a powerful psychological resource^22^. Nostalgic music, therefore, consists of familiar tunes that are strongly linked to an individual’s past and evoke mixed emotional experiences. Since music deeply connected to a person’s past can evoke these complex emotions, it may further strengthen autobiographical recall^14^. Thus, nostalgic music could be particularly effective in helping dementia patients with their recollections. Additionally, nostalgic reminiscence is known to enhance psychological resources by acting as a buffer against anxiety arising from perceived threats^23^. This effect of nostalgic music is especially significant in clinical settings, where it can serve as a valuable tool for dementia patients who often face existential threats due to their condition^24,25^.

However, the impact of nostalgic music varies from person to person, as music-induced nostalgia depends on personal characteristics, contextual factors like emotional experience with the music, memory states, and the interaction of these factors^26,27^. For example, music-evoked nostalgia is related to arousal and affective valence intensity, but the intensity of valence is modulated by mood state. Personalized music listening has also been recommended in conventional music therapy for dementia patients^28,29^. By not only playing an individual’s nostalgic songs but also quantitatively assessing the emotional state associated with each song during selection, it may be possible to maximize the feeling of nostalgia and make it more effective as a form of care for dementia patients.

To address individual differences, we developed a Nostalgia Brain-Music Interface (N-BMI), designed to enhance feelings of nostalgia by recommending songs tailored to each individual based on prediction models of nostalgic feelings. We tested this interface on healthy adults, which represents the first step toward future application in dementia patients. These models were built using rating data on subjective nostalgia, acoustic features of self-selected and other-selected nostalgic songs, and in-ear electroencephalographic (EEG) activity recorded while listening to the songs. The development of the N-BMI was inspired by previous research on EEG-based Brain-Computer Interfaces (BCIs). For instance, BCIs have been shown to modify music according to an individual’s estimated arousal and valence^30^ or to mediate emotions by generating personalized music^31^, highlighting the potential for recommending personalized music through EEG activity analysis. Additionally, a prior study demonstrated that neural activation related to nostalgia can be predicted by personality traits^32^, suggesting the feasibility of developing a neurofeedback system based on nostalgia-related brain activity. However, no research to date has developed a neurofeedback system specifically targeting nostalgia or examined its effects on nostalgia, well-being, and memory retrieval. Therefore, our goal was to develop the N-BMI and evaluate whether the songs recommended by the N-BMI could enhance nostalgic feelings, well-being, and memory recall in both young and elderly individuals.

## Methods

### Nostalgia Brain-Music Interface (N-BMI)

#### Overview

The proposed Nostalgia Brain-Music Interface (N-BMI) consisted of three steps, referred to as ‘Rec-Dec-Back’: (1) recording, (2) decoding, and (3) feedback (**Figure 1**). Before the experiment, participants selected three nostalgic songs. During the recording step, they listened to both self-selected and other-selected songs and rated their subjective feelings of nostalgia, well-being, and memory vividness. EEG activity was recorded using an in-ear EEG device (VIE Inc., Kanagawa, Japan) while participants listened to both self-selected and other-selected songs.

**Figure 1.**
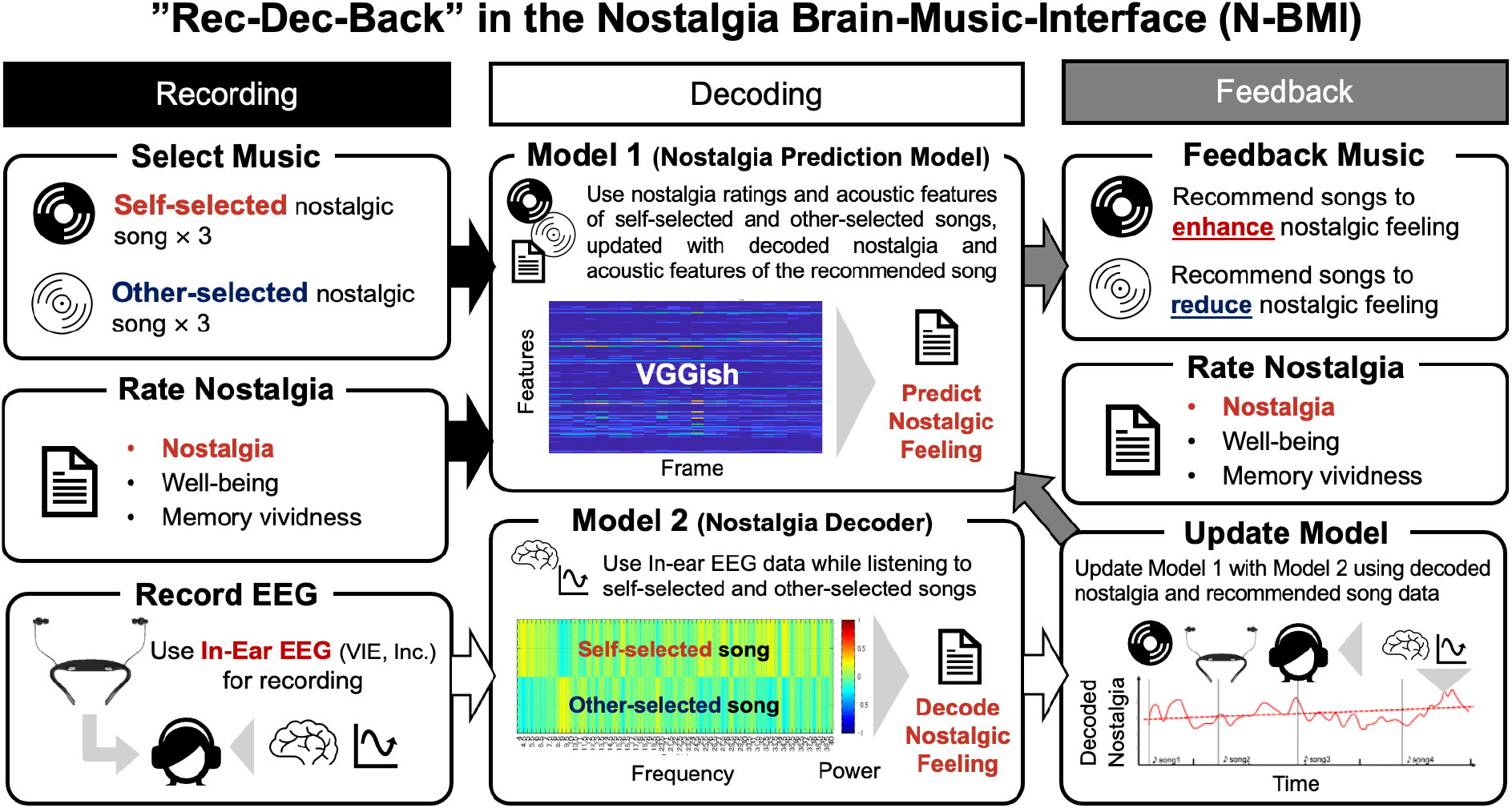
‘Rec-Dec-Back’ in the Nostalgia Brain-Music Interface (N-BMI). The N-BMI followed a three-step process called ‘Rec-Dec-Back,’ which refers to (1) recording, (2) decoding, and (3) feedback. During the recording step, participants listened to both self-selected and other-selected nostalgic songs and rated their subjective feelings of nostalgia, well-being, and memory vividness, while EEG activity was simultaneously recorded using an in-ear EEG device. In the decoding step, two models were developed: Model 1, the ‘Nostalgia Prediction Model,’ which utilized nostalgia ratings and the acoustic features of self-selected and other-selected songs extracted through the VGGish model; and Model 2, the ‘Nostalgia Decoder,’ which analyzed Fourier-transformed EEG power-frequency data recorded during song listening. Model 2 decoded nostalgic feelings by estimating whether the EEG patterns more closely matched those recorded during self-selected or other-selected song listening. In the feedback step, Model 1 predicted the participant’s nostalgic feelings and recommended a song from a pool of 7,481 songs to either enhance or reduce nostalgia. While the participant listened to the recommended song, Model 2 decoded nostalgic feelings, and both the decoded nostalgia and the song’s acoustic features were used to update Model 1. The updated Model 1 then predicted nostalgic feelings again and recommended another song from the pool. After listening to the recommended songs, participants rated their nostalgic feelings, well-being, and memory vividness.

In the decoding step, two models were created: Model 1, the ‘Nostalgia Prediction Model,’ utilized nostalgia ratings and the acoustic features of self-selected and other-selected songs. The acoustic features were extracted using the VGGish model, a pretrained convolutional neural network developed by Google^33,34^.It has been shown that the CNN model’s embedding is more sensitive to emotional responses than low-level acoustic features, and we thought it would be appropriate for use in N-BMI^35^. Model 2, the ‘Nostalgia Decoder,’ employed Fourier-transformed multidimensional EEG power-frequency data recorded while participants listened to the self-selected and other-selected songs. This model decoded nostalgic feelings from the EEG data by estimating whether the EEG pattern was more similar to those recorded during self-selected song listening or other-selected song listening.

In the feedback step, Model 1 predicted the participant’s nostalgic feeling and recommended a song from a pool of 7,481 songs to either enhance or reduce their nostalgia. While the participant listened to the recommended song, Model 2 decoded their nostalgic feeling every 20 seconds. The decoded nostalgia and the acoustic features of the recommended song were then added as new supervised inputs to update Model 1. The updated Model 1 predicted the nostalgic feeling again and recommended another song from the pool. During the feedback step, each song was played for 20 seconds, the model was updated every 20 seconds, and a total of six songs were played over 120 seconds (two minutes). After listening to the recommended songs, participants rated their nostalgic feelings, well-being, and memory vividness (see Recording in the Nostalgia Brain-Machine-Interface (N-BMI) section).

#### EEG Data Acquisition and Processing

We used an in-ear EEG device developed by VIE Inc. (Kanagawa, Japan, https://www.vie.style/), which records EEG data from the left and right ear canals using ear-tip-shaped electrodes, with reference electrodes placed at the back of the neck. The system records data from the left and right ear channels, as well as the difference signal between the two channels, to exclude noise from the common reference electrode signals. The ear-tip design allows for simultaneous recording of EEG signals and music playback. A previous study confirmed the similarity between in-ear EEG signals and temporal EEG signals measured by a conventional EEG device (Brain Amp DC, BrainVision, Canada)^36^.

The in-ear EEG data were recorded at a sampling frequency of 600 Hz and filtered using a fourth-order Butterworth filter with a frequency range of 3 to 40 Hz. The recorded data from the left and right channels, along with the difference-signal data, were segmented into 4-second time windows with 50% overlap. A Fourier transform was applied to each segment for each time-series data (i.e., left, right, and difference signal). EEG power across the 4-40 Hz bands was calculated in 0.5 Hz steps.

To detect noise, we performed two steps. First, to remove common noise between the two channels and over time, such as noise from eye movements, we applied independent component analysis (ICA) to the EEG data from the left and right channels. One of the components, which had the larger root mean square (RMS) value of the two components, was labeled as noise and removed from the filtered data. Second, to remove independent noise from each channel or time segment, we calculated a noise flag for each segment. This flag was estimated by taking the average value of the sum of the power spectral density, RMS, maximum gradient, and kurtosis, each of which was transformed to a z-score. The resulting mean value was considered an absolute amplitude value. If this amplitude value was larger than 2.5, the noise flag was set to 1; if smaller, the noise flag was set to 0. This calculation was applied separately to the right and left channel data. We used only the epochs without noise flags. Noise quantification and rejection were performed for each segment based on criteria from a previous study^37^.

For the extraction of the EEG components, the power-transformed data from the left and right ear channels, along with the difference signal, were combined horizontally and compressed to a maximum of 150 dimensions using principal component analysis (PCA). If the number of these dimensions was less than 150, we used all the components. These dimensional data were then used to create Model 2, the ‘Nostalgia Decoder,’ which decoded the participants’ nostalgic state.

#### Recording in the Nostalgia Brain-Machine-Interface (N-BMI)

Young participants were instructed to bring three songs that made them feel nostalgic prior to the experiment. Elderly participants were asked to choose from a list of songs that were popular when they were 15 years old, a period commonly associated with heightened autobiographical salience^10^. The elderly participants selected from this list because they were considered to have more difficulty recalling their nostalgic songs.

Participants wore in-ear EEG devices. During the recording step of the N-BMI, they listened to three self-selected nostalgic songs and three other-selected songs (six songs in total). Each song was played for 45 seconds from the beginning, with the order of the songs randomized. EEG data were recorded while participants listened to the six songs. After each song, participants rated their nostalgic feelings, well-being, and memory vividness using a Visual Analog Scale (VAS) ranging from 0 to 100 (0 = Not at all, 100 = Strongly). The VAS was displayed on a personal computer (PC), and participants were asked to enter their ratings. For elderly participants who had difficulty using the PC, a printed version of the VAS was provided, and they indicated their rating by pointing to the desired value on the paper, which the experimenter then inputted into the PC.

For the rating of nostalgic feelings, we used one item: (i) ‘I felt nostalgic,’ which participants answered on the VAS. For the rating of well-being, we used eight items: (i) ‘I was interested,’ (ii) ‘I felt loved and needed by others,’ (iii) ‘I had proper control over my actions, thoughts, and feelings,’ (iv) ‘I felt calm and peaceful,’ (v) ‘I was emotionally calm,’ (vi) ‘I was relaxed without problems,’ (vii) ‘I was cheerful and lighthearted,’ and (viii) ‘I was a happy person.’ These items were derived from an abbreviated version of the 38-item Mental Health Inventory (MHI-18)^38^. For the rating of memory vividness, we used three items: (i) ‘The overall memory of the event is extremely clear,’ (ii) ‘The memory of the event is extremely detailed,’ and (iii) ‘Overall, I remember the event clearly.’ These items were retrieved from the Japanese version of the Memory Characteristics Questionnaire (MCQ)^39^.

For elderly participants who used a printed version of the VAS, a shorter version of the well-being and memory vividness scales was administered to reduce the burden of answering multiple items. Specifically, they answered three items for well-being: (i) ‘I was interested,’ (iv) ‘I felt calm and peaceful,’ and (vii) ‘I was cheerful and lighthearted,’ and one item for memory vividness: (iii) ‘Overall, I remember the event clearly.’ This modification was made to accommodate elderly participants who had difficulty using the PC.

#### Decoding in the Nostalgia Brain-Machine Interface (N-BMI)

##### Model 1: Nostalgia Prediction Model

Model 1 (the Nostalgia Prediction Model) consisted of a Least Absolute Shrinkage and Selection Operator (LASSO) regression model that predicted nostalgic feelings based on acoustic features. Model 1 was developed using nostalgia ratings and acoustic feature data collected during the recording phase, which values were standardized to z-scored. Acoustic features were calculated using the VGGish model, which transforms audio inputs into spectrograms and extracts 128 dimensions of acoustic features. Each of the six songs played had a duration of 45 seconds, and VGGish transformed this song data into a matrix with 128 acoustic feature dimensions as columns and 184 time frames as rows—the number of time frames being determined independently by VGGish based on the song’s duration. An average value was then calculated for every 10-second segment for each dimension. These features were subsequently z-scored using the mean and standard deviation of the acoustic features from the songs in our music database (7,481 songs used in the feedback phase of the N-BMI, see below). If the standard deviation of a normalized feature exceeded an absolute value of 2, it was capped at 2 or -2. To reduce dimensionality, only the normalized features with a standard deviation greater than 0.01 were retained, as features below this threshold were considered to provide minimal informational value. Each segment was paired with nostalgia ratings for the corresponding piece of music, and this data was used as training data for Model 1.

Thus, this nostalgia prediction model can be described as: (1)

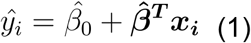

Where 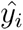 is the predicted nostalgic feeling for song *i*. 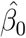 is the intercept, and 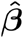 is a vector of coefficients for each acoustic feature. ***x***_*i*_ is a vector of acoustic features calculated by VGGish for song *i*, with values that have already been capped based on their standard deviation. 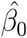 and 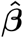 are the variables of *β*_0_ and *β*, which minimize the loss function expressed below:

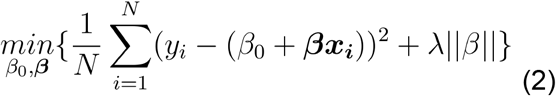

where λ is the optimal parameter to regulate overfitting, which is determined by cross validation (see Evaluation of Model Accuracy section). *N* represents the number of the training data, which is 6.

##### Model 2: Nostalgia Decoder

Model 2 (Nostalgia Decoder) consisted of a logistic LASSO regression model, which classified and calculated the likelihood that the current EEG pattern was closer to the pattern observed while listening to self-selected nostalgic songs or other-selected nostalgic songs. The model was created using the Fourier-transformed 150-dimensional EEG power-frequency data recorded during the listening sessions for both self-selected and other-selected songs (see EEG Data Acquisition and Processing section).

Consequently, this model can be described as:

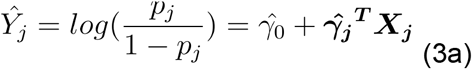

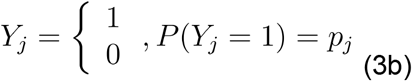

Where *Ŷ* represents the odds ratio of nostalgic feeling. *j* is the number of EEG frequency dimensions (with a maximum value of 150), 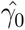 is the intercept, and 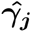 is a vector of coefficients corresponding to each EEG frequency. The optimal λ was selected by 20-fold cross-validation. These values were determined to minimize the loss function of below:

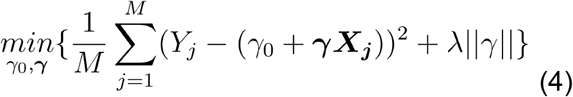

where λ is the optimal parameter to regulate overfitting, which is determined by cross validation (see Evaluation of Model Accuracy section). *M* represents the number of the training data, which is the number of the EEG frequency. The process was the same as used in Model 1. ***X***_*j*_ is a vector of dimensions representing EEG frequency. The model uses the EEG data in the most recent 4 seconds for decoding, which is performed every 1 second. After that, the decoded values for the last 5 seconds were averaged to smooth out sudden changes in decoding results. This moving mean value was used in the following analysis as the final decoded value. The methodology for creating the decoder model was the same as that used in the previous study^37^.

#### Feedback Using the Nostalgia Brain-Machine Interface (N-BMI)

##### Music Database Used for Feedback

We used a music database of 7,481 songs to generate music feedback in our N-BMI. The database consisted of songs that appeared in GfK Japan’s weekly music chart Top 1,000 between April 2018 and February 2023, as well as 600 songs popular from the 1920s to the 1980s, sourced from the “Seishun Uta Nenkan (Youth Song Yearbook)” CD collection.

##### Music Recommendation Using Models 1 and 2

First, Model 1 was applied to each song in the music database, and the predicted nostalgic feeling was calculated for each song. Next, we calculated the acoustic similarity (z-scored correlation coefficient) of each song in the music database relative to the averaged acoustic features of the three self-selected songs. We then summed the predicted nostalgic feeling and the acoustic similarity value to create a measure for music recommendation, which was used to rank the songs in the database. The higher the music recommendation value, the more similar the song was to the self-selected nostalgic songs and the more likely it was to induce a nostalgic feeling.

Using the music recommendation value, we recommended music under two conditions to test the effects of our N-BMI system: the “nostalgia condition,” where a song was recommended to enhance nostalgic feelings, and the “control condition,” where a song was recommended to reduce nostalgic feelings (see **Figures 2A and 2B**, respectively). In the nostalgia condition, the system randomly selected a song from the top 10 songs ranked by the music recommendation value, while in the control condition, the system randomly selected a song from the bottom 10 songs ranked by the music recommendation value.

**Figure 2.**
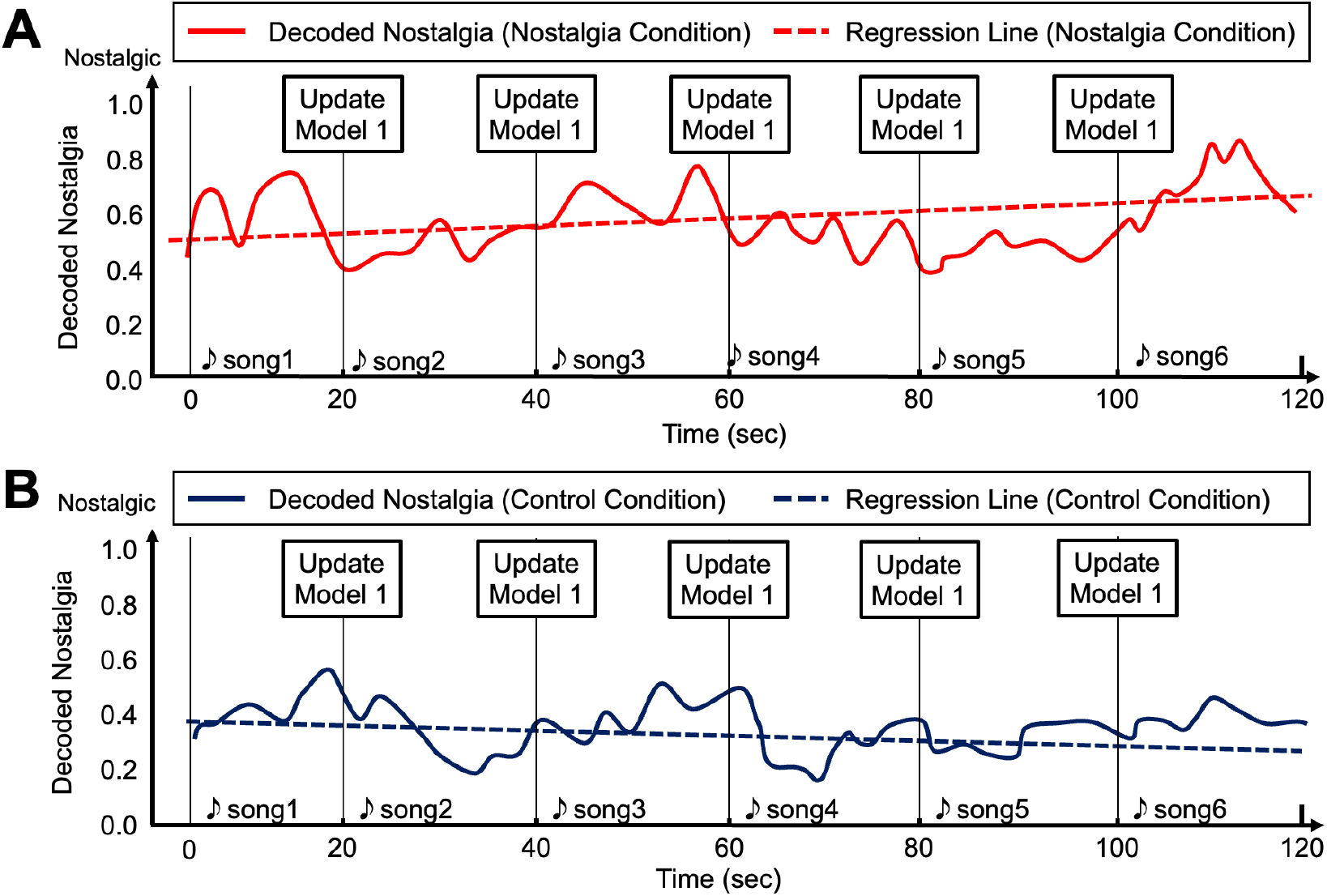
Schematic drawing of decoded nostalgia while listening to songs recommended by the Nostalgia Brain-Music Interface (N-BMI). The red solid and dashed lines represent the decoded nostalgia and regression line during the nostalgia condition, respectively, where the song was recommended to enhance nostalgic feelings (**A**). The blue solid and dashed lines represent the decoded nostalgia and regression line during the control condition, respectively, where the song was recommended to reduce nostalgic feelings (**B**). In both conditions, each recommended song was played for 20 seconds, and Model 1 was updated every 20 seconds after each song. A total of six songs were recommended over a 120-second period.

After the first recommended song was played for 20 seconds, Model 2 decoded the nostalgic feeling from in-ear EEG data (**Figure 2**). Using the decoded nostalgic-feeling value (*Ŷ* value in (3a)) and the maximum and minimum VAS values of each participant, we calculated the 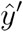 value as:

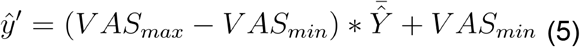

The 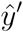 the decoded nostalgic feeling, which is the to approximate VAS value using the maximum and minimum VAS ratings of nostalgia in a participant. This approximate value 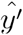 was used for updating after standardization to z-score. 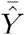 is the average of the decoded nostalgic feeling over the 20 seconds of data recording. The average of 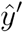 value and the VGGish values of the recommended song over the 20 seconds were used to update Model 1. Thus, Model 1 was updated every 20 seconds during the music feedback. Using the updated Model 1, our system recalculated the music recommendation value and re-ranked the songs in the database. During the music feedback step, (1) was updated five times in total and recommended six songs over 120 seconds (two minutes) (**Figure 2**).

### Experiment

#### Participants

Thirty-four participants took part in the experiment: seventeen elderly participants (age range: 70–87, M = 80.6, SD = 4.4 years; 3 male) from a Rehabilitation Day Care Center, and seventeen young participants (age range: 21–35, M = 26.8, SD = 7.1 years; 9 male). All participants were given both oral and written explanations of the experiment, and each provided written informed consent. The study was approved by the Ethical Committee of Shiba Palace Clinic (approval number: 152115_rn-34967).

The elderly participants completed the Mini-Cog test^40^, a simple dementia screening tool that includes a recall test (maximum score: 3) and a clock drawing test (maximum score: 2). The total Mini-Cog score ranges from 0 to 5, with elderly participants in this study scoring between 0 and 5 (M = 3.3, SD = 1.5 points). Since a total score of 0, 1, or 2 suggests a higher likelihood of clinically significant cognitive impairment, the elderly participants represented individuals with varying degrees of cognitive impairment.

#### Experimental Procedure

Both young and elderly participants selected three songs that evoked feelings of nostalgia. Participants also provided demographic information (age and gender). After providing their demographic data, the elderly participants completed the Mini-Cog test. The experimenter then moistened and cleaned the participants’ ears and the back of their necks, where electrodes were attached, before placing the in-ear EEG device.

Participants then listened to six songs—three self-selected and three chosen by others—in random order (“Recording” in **Figure 3**). After listening to each song, they rated their nostalgic feelings, well-being, and memory vividness (see details above). In-ear EEG signals were recorded while participants listened to the songs. Using this data, Models 1 and 2 were generated and their accuracy for each participant was evaluated (“Decoding” in **Figure 3**). Models 1 and 2 then recommended six songs for two conditions: the “nostalgia condition,” where songs were selected to enhance nostalgic feelings, and the “control condition,” where songs were chosen to reduce nostalgic feelings (“Feedback” in **Figure 3**). After listening to the six songs in each condition, participants again rated their nostalgic feelings, well-being, and memory vividness.

**Figure 3.**
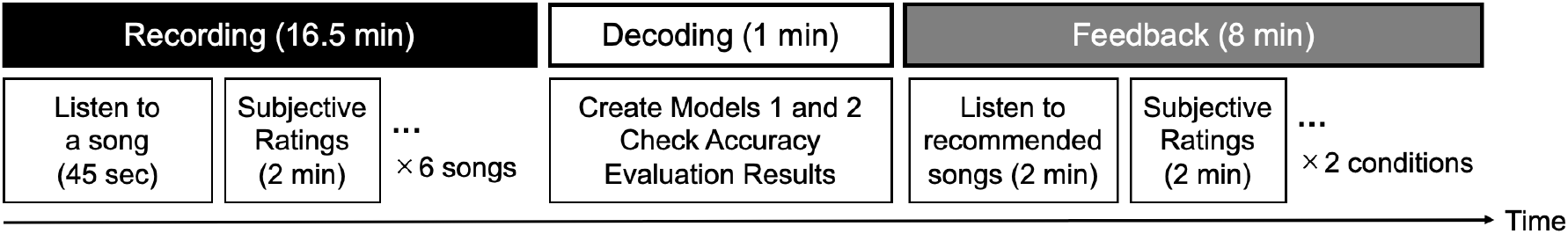
Experimental Procedure. Participants listened to six songs—three self-selected and three other-selected nostalgic songs—in random order (“Recording”). After listening to each song, they subjectively rated their nostalgic feelings, well-being, and memory vividness. Using the recording data, Models 1 and 2 were created and their accuracy was evaluated for each participant (“Decoding”). Based on these models, six songs were recommended for two conditions: the “nostalgia” condition, where songs were selected to enhance nostalgic feelings, and the “control” condition, where songs were selected to reduce nostalgic feelings (“Feedback”). After each condition, participants again subjectively rated their nostalgic feelings, well-being, and memory vividness.

### Analyses

#### Subjective Ratings During Recording

The subjective ratings during the recording step were analyzed as follows: First, the VAS ratings obtained for each item were averaged to measure nostalgic feelings, well-being, and memory vividness. Second, the averaged VAS ratings after listening to both self-selected and other-selected songs were calculated for each participant. Third, a Wilcoxon signed-rank test was conducted to compare the VAS ratings for self-selected versus other-selected songs for both the young and elderly participant groups.

#### Evaluation of Model Accuracy

For Model 1 (Nostalgia Prediction Model), the correlation coefficient between predicted and measured values after training was used as a measure of model accuracy. For Model 2 (Nostalgia Decoder), the model was trained on 90% of randomly selected EEG data windows (4-second windows with 50% overlap), and classification accuracy was calculated for the remaining 10% of the data. This was a binary classification task: self-selected nostalgic song or other-selected nostalgic song, with a chance level of 50%. During cross-validation, the optimal lambda (λ) value was chosen based on the lowest testing error. Classification accuracy was calculated for each participant and averaged across both the young and elderly participant groups. One-sample t-tests were conducted to determine whether the decoding accuracies for the young and elderly groups were significantly higher than the chance level of 50%. Additionally, the Area Under the Curve (AUC) was calculated for each participant and averaged for both groups.

#### Decoded Nostalgia and Subjective Ratings During Feedback

We calculated the mean decoded nostalgia over the entire 120-second period and the mean decoded nostalgia during the final 20 seconds for both the nostalgia and control conditions for each participant. Missing values were excluded when calculating the average value. The mean decoded nostalgia values over the 120-second period and during the last 20 seconds were compared between the nostalgia and control conditions using Wilcoxon signed-rank tests for both the young and elderly participant groups.

The subjective ratings during the feedback phase were analyzed as follows: First, the VAS ratings for each item were averaged to measure nostalgic feelings, well-being, and memory vividness. Second, the averaged VAS ratings after listening to the six songs in the nostalgia and control conditions were calculated for each participant. Third, a Wilcoxon signed-rank test was conducted to compare the average VAS ratings for songs in the nostalgia versus control conditions for both the young and elderly participant groups. All data analyses were performed using R version 4.3.3 and MATLAB 2023b software. A significance level of *p* < 0.05 was used for all statistical analyses. Wilcoxon signed-rank tests were employed for variables that did not meet the assumptions for parametric tests, such as normal distribution and/or homogeneity of variances, as assessed by the Shapiro-Wilk and Levene’s tests.

## Results

### Subjective Ratings During Recording

For the young participants, VAS ratings for nostalgic feelings, well-being, and memory vividness were significantly higher after listening to self-selected songs compared to other-selected songs (nostalgic feelings: p < 0.001, Cohen’s r = 0.878; well-being: p = 0.001, Cohen’s r = 0.741; memory vividness: p < 0.001, Cohen’s r = 0.878; see **Figure 4A**).

**Figure 4.**
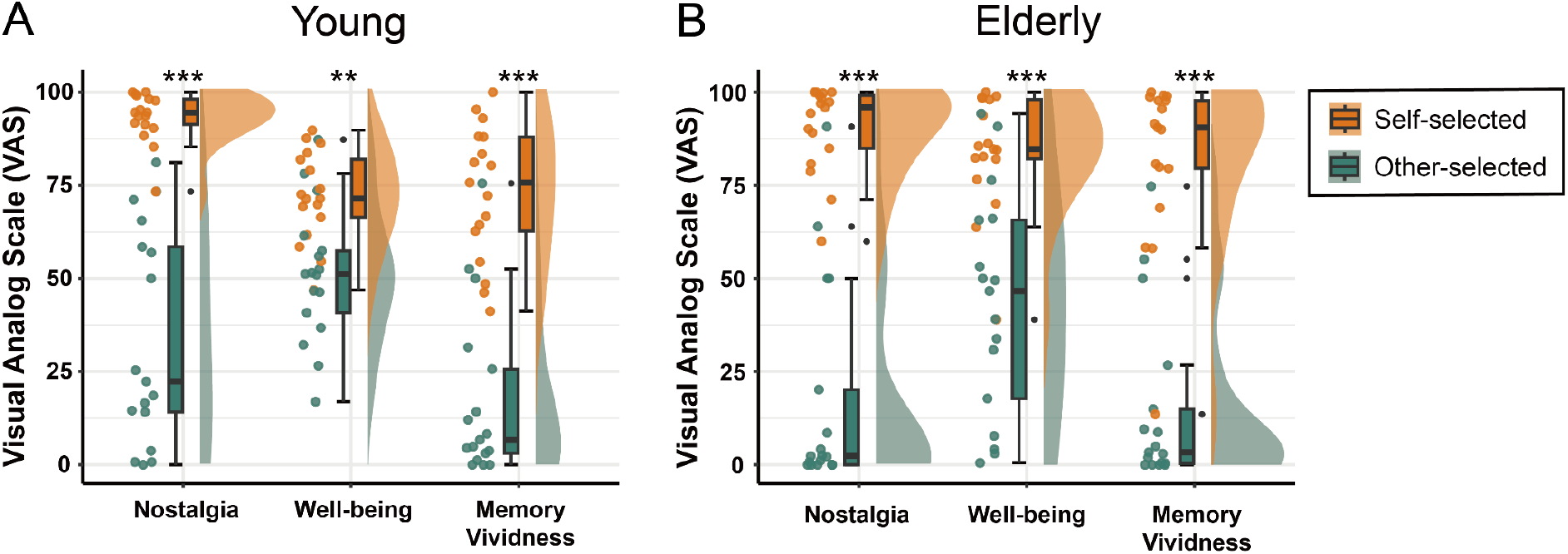
Subjective Visual Analog Scale (VAS) ratings for nostalgia feelings, well-being, and memory vividness after listening to self-selected and other-selected nostalgic songs. VAS ratings for young and elderly participants are shown in panels (A) and (B), respectively. Orange represents the data after listening to self-selected songs, while green represents the data after listening to other-selected songs. Each dot corresponds to an individual participant’s data. Error bars indicate the median ± 1.5 times the interquartile range (IQR). **p < .01, ***p < .001.

For the elderly participants, VAS ratings for nostalgic feelings, well-being, and memory vividness were also significantly higher after listening to self-selected songs compared to other-selected songs (nostalgic feelings: p < 0.001, Cohen’s r = 0.878; well-being: p < 0.001, Cohen’s r = 0.844; memory vividness: p < 0.001, Cohen’s r = 0.878; see **Figure 4B**).

#### Accuracy of Models 1 and 2

For Model 1, the mean correlation coefficients were high for both young participants (r = 0.985) and elderly participants (r = 0.995). For Model 2, the mean classification accuracy of the nostalgia decoder was 63.97 ± 16.11 (Mean ± Standard Deviation: SD) for young participants and 71.52 ± 19.88 (Mean ± SD) for elderly participants. One-sample t-tests showed that the mean classification accuracy of the nostalgia decoder was significantly higher than the chance level (50%) in both groups (Young: t(16) = 16.37, p < 0.01; Elderly: t(16) = 14.83, p < 0.01). The mean AUCs for young and elderly participants were 0.70 and 0.81, respectively.

#### Decoded Nostalgia During Feedback

The mean decoded nostalgia over the 120-second period and during the last 20 seconds in the nostalgia condition was not significantly higher than in the control condition for the young participants (p = 0.07, Cohen’s r = 0.362, **Figure 5A**). In contrast, while the mean decoded nostalgia during the last 20 seconds in the nostalgia condition was not significantly higher than in the control condition, but over the 120 seconds was significantly higher than in the control condition for the elderly participants (p = 0.003, Cohen’s r = 0.683, **Figure 5B**).

**Figure 5.**
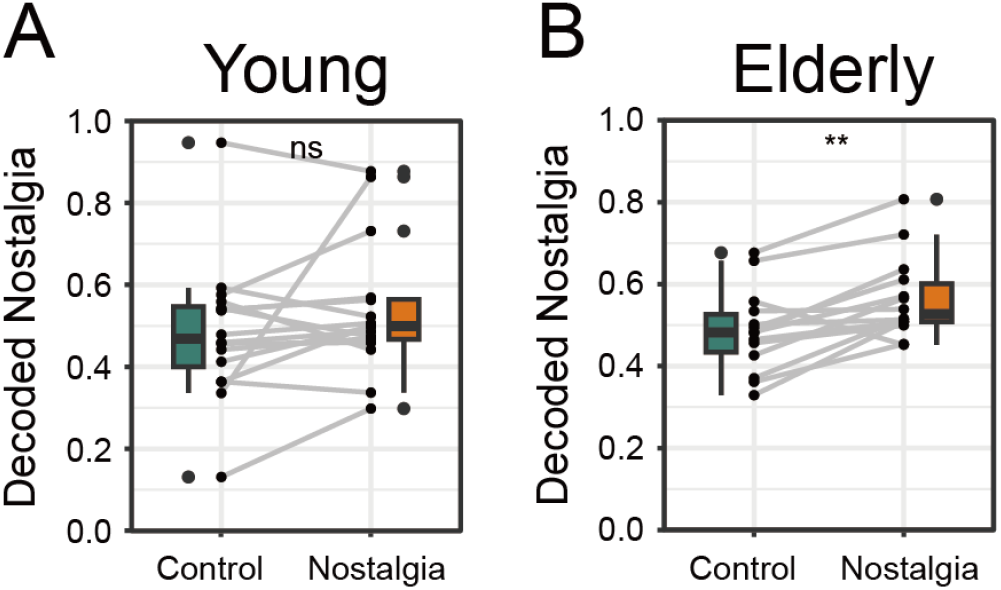
Decoded nostalgia estimated by Model 2 while listening to songs recommended by the Nostalgia Brain-Music Interface (N-BMI). (**A**) and (**B**) show the mean decoded nostalgia over 120 seconds for the control (green) and nostalgia (orange) conditions in young (**A**) and elderly (**B**) participants, respectively. Error bars represent the median ± 1.5 * IQR (interquartile range). n.s. = not significant. *p* < .05, \**p* < .01, \*\**p* < .001.

#### Subjective Ratings During Feedback

For the young participants, VAS ratings for nostalgic feelings, well-being, and memory vividness were significantly higher after listening to the recommended songs in the nostalgia condition compared to the control condition (nostalgic feelings: p = 0.003, Cohen’s r = 0.718; well-being: p = 0.003, Cohen’s r = 0.803; memory vividness: p = 0.036, Cohen’s r = 0.514; see **Figure 6A**).

**Figure 6.**
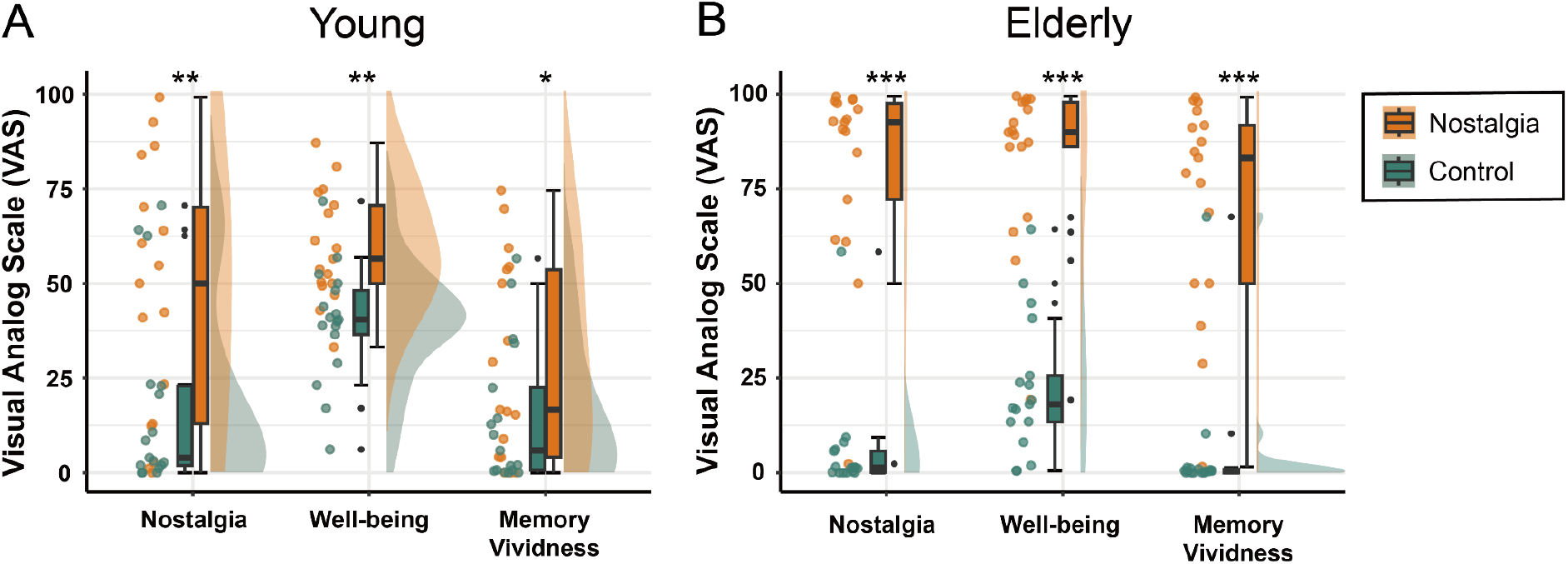
Subjective Visual Analog Scale (VAS) ratings for nostalgic feelings, well-being, and memory vividness after listening to songs recommended by the Nostalgia Brain-Music Interface (N-BMI). VAS ratings for young and elderly participants are shown in panels (A) and (B), respectively. Orange represents the data after listening to the song recommended to enhance nostalgic feelings (“nostalgia condition”), while green represents the data after listening to the song recommended to reduce nostalgic feelings (“control condition”). Each dot corresponds to data from an individual participant. Error bars represent the median ± 1.5 times the interquartile range (IQR). p < .05, *p < .01, **p < .001.

For the elderly participants, VAS ratings for nostalgic feelings, well-being, and memory vividness were also significantly higher after listening to the recommended songs in the nostalgia condition compared to the control condition (nostalgic feelings: p < 0.001, Cohen’s r = 0.878; well-being: p < 0.001, Cohen’s r = 0.878; memory vividness: p < 0.001, Cohen’s r = 0.878; see **Figure 6B**).

## Discussion

The aim of this study was to develop the Nostalgia Brain-Music Interface (N-BMI) and evaluate its ability to enhance nostalgic feelings, well-being, and memory recall in both young and elderly individuals. The N-BMI followed a three-step process called “Rec-Dec-Back,” which represents (1) recording, (2) decoding, and (3) feedback. In the recording phase, participants listened to both self-selected and other-selected nostalgic songs, rating their nostalgic feelings using the Visual Analog Scale (VAS), while in-ear EEG data were recorded throughout the session. In the decoding phase, models predicting participants’ nostalgic feelings were developed using VAS ratings, auditory features, and in-ear EEG data collected during the recording phase. In the feedback phase, the N-BMI recommended songs designed to either enhance or reduce nostalgic feelings (nostalgia vs. control conditions).

Our results demonstrated that VAS ratings of nostalgic feelings, well-being, and memory vividness were significantly higher in the nostalgia condition compared to the control condition for both young and elderly participants. These findings indicate that the N-BMI effectively enhanced nostalgic feelings, well-being, and memory recall across both age groups.

### Utilizing Self-Selected Songs to Enhance and Decode Nostalgia

We demonstrated that VAS ratings for nostalgic feelings, well-being, and memory vividness were significantly higher after listening to self-selected nostalgic songs compared to other-selected songs in both young and elderly participants (**Figure 4**), highlighting the importance of considering individual differences when evoking nostalgic feelings. This finding supports the idea that music-induced nostalgia varies from person to person^26,27^.

We propose that self-selected nostalgic songs were more closely tied to personal past experiences, inducing greater autobiographical salience compared to other-selected songs. This is consistent with the previous study^15^, which demonstrated that self-selected music more effectively enhanced autobiographical recall in patients with Alzheimer’s disease compared to researcher-selected music. Our findings further support the idea that nostalgic feelings are associated with autobiographical prominence^10,27^. Therefore, we suggest that self-selected nostalgic songs, being deeply connected to personal past experiences, induced greater autobiographical salience and resulted in more vivid autobiographical memories.

We also found that self-selected nostalgic songs enhanced subjective well-being more than other-selected songs. Previous studies have shown that music-evoked nostalgia, accompanied by autobiographical memories, enhances social bonding^41^, perceived meaning in life^42^, and self-esteem^43^. Our findings align with these studies, suggesting that music-evoked nostalgia enhances subjective well-being. This is also consistent with the idea that self-selected songs can be effective in clinical settings for supporting memory and well-being in patients with amnesia and dementia^15,44^.

Using in-ear EEG data recorded while participants listened to self-selected and other-selected songs, we developed Model 2 (Nostalgia Decoder), which predicts whether an EEG pattern is more similar to those recorded during self-selected or other-selected song listening. We demonstrated that the classification accuracy of Model 2 was significantly higher than chance level in both young and elderly participants. Thus, we suggest that comparing self-selected and other-selected songs is an effective approach for developing an EEG decoder of nostalgic feelings.

#### The Potential of the Nostalgia Brain-Music Interface (N-BMI) to Evoke Nostalgia, Enhance Memory, and Improve Well-Being

It is noteworthy that the N-BMI in this study evoked nostalgic feelings by recommending songs from a music database, even though these songs were not necessarily tied to the individual’s direct past experiences. It is possible that songs without prior listening experience evoked nostalgic feelings when recommended by the N-BMI.

Typically, nostalgia is triggered by stimuli that relate to past events and personal experiences. Previous research highlights that familiarity and autobiographical salience are key factors in the intensity of nostalgia^27^. Therefore, it was expected that the self-selected nostalgic songs in the recording phase of this study—songs directly experienced by the individuals and tied to personal past experiences—would evoke stronger nostalgic feelings. However, the songs recommended by the N-BMI were not necessarily tied to the individual’s direct past experiences. If so, why and how did the N-BMI evoke nostalgic feelings?

We propose that the dual-process model of recognition memory may explain this phenomenon. The dual-process model posits that two distinct cognitive processes, “recollection” and “familiarity,” are involved in recognition judgments^45,46^. Recollection involves retrieving specific details about a past event, while familiarity refers to a general sense that something has been encountered before, without recalling specific details^47^. Recollection operates through a threshold process, meaning it either occurs fully or not at all^46^, and involves conscious retrieval of past events. In contrast, familiarity is shaped by how well the features of a stimulus match previous experiences, with memory traces strengthening gradually through accumulated exposure^48^. This familiarity-based recognition, where the source memory is not accessible but a sense of recognition is felt, is thought to share mechanisms with déjà vu—the sensation of having experienced a current situation before, despite knowing it is novel^49^. An auditory version of déjà vu, known as déjà entendu, has been documented^50^, suggesting that novel sounds not directly tied to an individual’s past can evoke a sense of familiarity, triggering nostalgic feelings.

Considering the dual-process model and the déjà entendu phenomenon, we suggest that the songs recommended by the N-BMI may evoke nostalgic feelings and enhance memory vividness primarily through the familiarity process rather than recollection. Since Model 1 in our N-BMI algorithm recommended songs based on acoustic feature similarity, the features of recommended songs matched those of self-selected nostalgic songs that were tied to past experiences. This matching could have increased the sense of accumulated familiarity, thereby evoking nostalgic feelings and enhancing memory vividness. Additionally, previous studies have shown that music aids in retaining detailed memories^51^ and encoding context, which particularly enhances verbal episodic memory^52^. The recommended songs, with similar acoustic features to participants’ nostalgic songs, may have helped participants access more detailed past memories.

The N-BMI not only enhanced memory vividness but also improved well-being in both young and elderly participants. These results are consistent with previous research highlighting the role of music and nostalgia in promoting psychological health^22,53^. We propose that the N-BMI improved well-being through similar mechanisms: enhanced feelings of nostalgia and strengthened memory recall could evoke positive emotions, reinforce social bonds, increase self-esteem, and reduce negative affect, ultimately improving well-being.

Taken together, we suggest that the N-BMI has the potential to evoke nostalgia, enhance memory, and improve well-being by recommending songs from a database, even when participants had no prior listening experience with those songs.

#### Age-Related Differences in Enhancing and Decoding Nostalgia

We found that VAS ratings for nostalgic feelings, well-being, and memory vividness were significantly higher after listening to the recommended songs in the nostalgia condition compared to the control condition for both young and elderly participants (**Figure 6**). However, effect sizes for elderly participants (Cohen’s r = 0.878) were larger than those for younger participants (Cohen’s r = 0.514–0.803). Thus, although the N-BMI was effective in enhancing subjective nostalgic feelings, well-being, and memory vividness in both age groups, the effect size was greater for the elderly population than for the younger population.

This age-related difference was also reflected in the accuracy of Model 2, the Nostalgia Decoder. Specifically, the mean decoded nostalgia was significantly higher in the nostalgia condition than in the control condition for elderly participants (**Figures 5B**), suggesting that the EEG pattern during the nostalgia condition more closely aligned with that observed while listening to self-selected nostalgic songs. In contrast, there was no significant difference in the mean decoded nostalgia between the two conditions (**Figures 5A**). Thus, the age-related differences in the decoded nostalgia results correspond to those observed in the VAS ratings.

We propose at least three factors to explain these age-related differences. First, variation in acoustic features between older and more recent songs in our music database may have influenced the results. Our database included popular Japanese songs from 2018 to 2023, likely familiar to younger participants, and songs from the 1920s to 1980s, familiar to elderly participants. Since songs from different eras possess distinct acoustic features, this likely affected the music recommendation process. For elderly participants, older songs from their youth were ranked higher, while recent songs were ranked lower (**Figure 4B)**, leading to the recommendation of recent songs in the control condition. It is plausible that elderly participants did not experience nostalgia in the control condition because recent songs are less likely to evoke nostalgia. Conversely, for younger participants, recent songs were ranked higher, while older songs were ranked lower, leading to the recommendation of older songs in the control condition. Younger participants may have still experienced nostalgia in the control condition, as older songs could be linked to cultural or collective memory. For instance, younger participants familiar with nostalgic stories related to older songs may have experienced nostalgia from those songs, diminishing the difference between the nostalgia and control conditions.

Second, age-related differences in response style bias may also explain the results. Aging and cognitive decline are known to lead to more extreme response styles^54^. Consistent with this notion, elderly participants in this study exhibited more extreme responses on the VAS (**Figures 4 and 6**). Since VAS data collected during the recording phase was used to create Model 1 (the Nostalgia Prediction Model), this response bias may have contributed to the relatively higher accuracy of the model for elderly participants (r = 0.995) compared to younger participants (r = 0.985). This bias may also explain the more pronounced differences in VAS ratings during the feedback phase.

Third, age-related differences in the vividness of memory recall may have influenced the results. Previous research has shown that elderly participants tend to recall memories more vividly^55^. If this is the case, elderly participants may have benefited more from the N-BMI simply because they recalled memories more vividly, likely due to stronger and more detailed connections to past events.

#### Future Directions

A potential future direction is applying the N-BMI to patients with dementia and Alzheimer’s disease. The results of this study suggest that the N-BMI could be particularly beneficial for patients with severe memory impairments, making it a promising area for future research. Additionally, the N-BMI holds potential as a non-pharmacological approach to dementia prevention, offering a feasible, low-cost, and low-side-effect option that can be implemented at home. The findings from this study suggest that this drug-free dementia prevention method could be integrated into home settings.

Another future direction is to apply the N-BMI in broader contexts. Since the N-BMI offers a new way of recommending music using an in-ear EEG device, it could be easily adapted for use in the general population in daily life. It has the potential to enhance nostalgia, memory vividness, and well-being in individuals across various demographics. Future studies could explore its impact on emotional health in diverse populations. The development of such technologies could provide significant benefits for individuals with memory and emotional regulation challenges, paving the way for innovative therapeutic interventions in a wider range of contexts.

## Conclusion

We developed the Nostalgia Brain-Music Interface (N-BMI) and demonstrated that it enhanced nostalgic feelings, well-being, and memory recall in both young and elderly individuals. The N-BMI paves the way for innovative therapeutic interventions, including a non-pharmacological approach for patients with dementia, across a wider range of contexts.

## Acknowledgements

We would like to thank Dr. Seung-goo Kim for his warm advice on the study.

## Author contributions

YS: conceptualization, data curation, formal analysis, methodology, visualization, writing – original draft, writing – review & editing, validation. TK: conceptualization, methodology, data curation, writing – review & editing. SK: writing – review & editing. TE: writing – review & editing. SS: data curation, writing – review & editing. YI: conceptualization, writing – review & editing. YN: conceptualization, writing – review & editing. SF: conceptualization, software, methodology, writing – original draft, writing – review & editing. TI: conceptualization, data curation, formal analysis, visualization, methodology, software, writing – review & editing.

## Data availability statement

The datasets analyzed in the current study are available in the Open Science Frameworks (OSF) repository, https://osf.io/yfcuk/.

## Reference

1. Sheehan, B. Assessment scales in dementia. Ther. Adv. Neurol. Disord. 5, 349–358. (2012).

2. GBD 2019 Dementia Forecasting Collaborators. Estimation of the global prevalence of dementia in 2019 and forecasted prevalence in 2050: an analysis for the Global Burden of Disease Study 2019. Lancet Public Health 7, e105–e125 (2022).

3. Zucchella, C. et al. The multidisciplinary approach to Alzheimer’s disease and dementia. A narrative review of non-pharmacological treatment. Front. Neurol. 9, 1058 (2018).

4. Li, Y.-Q. et al. Non-pharmacological interventions for behavioral and psychological symptoms of dementia: A systematic review and network meta-analysis protocol. Front. Psychiatry 13, 1039752 (2022).

5. Matziorinis, A. M. & Koelsch, S. The promise of music therapy for Alzheimer’s disease: A review. Ann. N. Y. Acad. Sci. 1516, 11–17 (2022).

6. Petersen, R. C. et al. Current concepts in mild cognitive impairment. Arch. Neurol. 58, 1985–1992 (2001).

7. American Psychiatric Association. Diagnostic and Statistical Manual of Mental Disorders: DSM-5-TR.American Psychiatric Association Publishing, 2022).

8. Ménard, M.-C. & Belleville, S. Musical and verbal memory in Alzheimer’s disease: a study of long-term and short-term memory. Brain Cogn. 71, 38–45 (2009).

9. Jacobsen, J.-H. et al. Why musical memory can be preserved in advanced Alzheimer’s disease. Brain 138, 2438–2450 (2015).

10. Janata, P., Tomic, S. T. & Rakowski, S. K. Characterisation of music-evoked autobiographical memories. Memory 15, 845–860 (2007).

11. Foster, N. A. & Valentine, E. R. The effect of auditory stimulation on autobiographical recall in dementia. Exp. Aging Res. 27, 215–228 (2001).

12. Meilán García, J. J. et al. Improvement of autobiographic memory recovery by means of sad music in Alzheimer’s Disease type dementia. Aging Clin. Exp. Res. 24, 227–232 (2012).

13. Fraile, E. et al. The effect of learning an individualized song on autobiographical memory recall in individuals with Alzheimer’s disease: A pilot study. J. Clin. Exp. Neuropsychol. 41, 760–768 (2019).

14. El Haj, M., Fasotti, L. & Allain, P. The involuntary nature of music-evoked autobiographical memories in Alzheimer’s disease. Conscious. Cogn. 21, 238–246 (2012).

15. El Haj, M., Antoine, P., Nandrino, J. L., Gély-Nargeot, M.-C. & Raffard, S. Self-defining memories during exposure to music in Alzheimer’s disease. Int. Psychogeriatr. 27, 1719–1730 (2015).

16. Cuddy, L. L., Sikka, R. & Vanstone, A. Preservation of musical memory and engagement in healthy aging and Alzheimer’s disease: Musical memory in Alzheimer’s disease. Ann. N. Y. Acad. Sci. 1337, 223–231 (2015).

17. Moreira, S. V., Justi, F. R. D.R. & Moreira, M. Can musical intervention improve memory in Alzheimer’s patients? Evidence from a systematic review. Dement Neuropsychol 12, 133–142 (2018).

18. Bierman, E. J. M., Comijs, H. C., Jonker, C. & Beekman, A. T. F. Symptoms of anxiety and depression in the course of cognitive decline. Dement. Geriatr. Cogn. Disord. 24, 213–219 (2007).

19. Ueda, T., Suzukamo, Y., Sato, M. & Izumi, S.-I. Effects of music therapy on behavioral and psychological symptoms of dementia: a systematic review and meta-analysis. Ageing Res. Rev. 12, 628–641 (2013).

20. Raglio, A. et al. Efficacy of music therapy in the treatment of behavioral and psychiatric symptoms of dementia. Alzheimer Dis. Assoc. Disord. 22, 158–162 (2008).

21. Särkämö, T. et al. Music, emotion and dementia: Insight from neuroscientific and clinical research. Music and Medicine 4, 153–162 (2012).

22. Sedikides, C., Rudich, E. A., Gregg, A. P., Kumashiro, M. & Rusbult, C. Are normal narcissists psychologically healthy?: self-esteem matters. J. Pers. Soc. Psychol. 87, 400–416 (2004).

23. Sedikides, C., Wildschut, T., Routledge, C. & Arndt, J. Nostalgia counteracts self-discontinuity and restores self-continuity. Eur. J. Soc. Psychol. 45, 52–61 (2015).

24. Umar Ismail, S., Cheston, R., Christopher, G. & Meyrick, J. Nostalgia as a psychological resource for people with dementia: A systematic review and meta-analysis of evidence of effectiveness from experimental studies. Dementia 19, 330–351 (2020).

25. Ismail, S. et al. Psychological and Mnemonic Benefits of Nostalgia for People with Dementia. J. Alzheimers. Dis. 65, 1327–1344 (2018).

26. Hennessy, S., Greer, T., Narayanan, S. & Habibi, A. Unique affective profile of music-evoked nostalgia: An extension and conceptual replication of Barrett et al.’s (2010) study. Emotion (2024) doi:10.1037/emo0001389.

27. Barrett, F. S. et al. Music-evoked nostalgia: affect, memory, and personality. Emotion 10, 390–403 (2010).

28. Rao, C. B., Peatfield, J. C., McAdam, K. P. W. J., Nunn, A. J. & Georgieva, D. P. A focus on the reminiscence bump to personalize music playlists for dementia. J. Multidiscip. Healthc. 14, 2195–2204 (2021).

29. Murphy, K. et al. Implementation of personalized music listening for assisted living residents with dementia. Geriatr. Nurs. 39, 560–565 (2018).

30. Ramirez, R., Palencia-Lefler, M., Giraldo, S. & Vamvakousis, Z. Musical neurofeedback for treating depression in elderly people. Front. Neurosci. 9, 354 (2015).

31. Ehrlich, S. K., Agres, K. R., Guan, C. & Cheng, G. A closed-loop, music-based brain-computer interface for emotion mediation. PLoS One 14, e0213516 (2019).

32. Barrett, F. S. & Janata, P. Neural responses to nostalgia-evoking music modeled by elements of dynamic musical structure and individual differences in affective traits. Neuropsychologia 91, 234–246 (2016).

33. Gemmeke, J. F. et al. Audio Set: An ontology and human-labeled dataset for audio events. in 2017 IEEE International Conference on Acoustics, Speech and Signal Processing (ICASSP) 776–780 (IEEE, 2017). doi:10.1109/ICASSP.2017.7952261.

34. Hershey, S. et al. CNN architectures for large-scale audio classification. in 2017 IEEE International Conference on Acoustics, Speech and Signal Processing (ICASSP) 131–135 (IEEE, 2017). doi:10.1109/ICASSP.2017.7952132.

35. Seung-Goo, K., Tobias, O. & Daniela, S. Emotion-relevant Representations of Music Extracted by Convolutional Neural Networks Are Encoded in Medial Prefrontal Cortex. Preprint at (2023).

36. Ueda, K., Imamura, Y. & Ibaraki, T. The development of a eustress sensing system using In-Ear EEG. in 2021 43rd Annual International Conference of the IEEE Engineering in Medicine & Biology Society (EMBC) (IEEE, 2021). doi:10.1109/ACII.2019.8925442.

37. Chang, M., Ibaraki, T., Naruse, Y. & Imamura, Y. A study on neural changes induced by sauna bathing: Neural basis of the ‘totonou’ state. PLoS One 18, e0294137 (2023).

38. Veit, C. T. & Ware, J. E. The structure of psychological distress and well-being in general populations. J. Consult. Clin. Psychol. 51, 730–742 (1983).

39. Takahashi, M. & Shimizu, H. Do you remember the day of your graduation ceremony from junior high school?: A factor structure of the Memory Characteristics Questionnaire1. Jpn. Psychol. Res. 49, 275–281 (2007).

40. Borson, S., Scanlan, J. M., Chen, P. & Ganguli, M. The Mini-Cog as a screen for dementia: validation in a population-based sample. J. Am. Geriatr. Soc. 51, 1451–1454 (2003).

41. Cheung, W.-Y. et al. Back to the future: nostalgia increases optimism. Pers. Soc. Psychol. Bull. 39, 1484–1496 (2013).

42. Routledge, C. et al. The past makes the present meaningful: nostalgia as an existential resource. J. Pers. Soc. Psychol. 101, 638–652 (2011).

43. Hart, C. M. et al. Nostalgic recollections of high and low narcissists. J. Res. Pers. 45, 238–242 (2011).

44. Arroyo-Anlló, E. M., Díaz, J. P. & Gil, R. Familiar music as an enhancer of self-consciousness in patients with Alzheimer’s disease. Biomed Res. Int. 2013, 752965 (2013).

45. Diana, R. A., Reder, L. M., Arndt, J. & Park, H. Models of recognition: a review of arguments in favor of a dual-process account. Psychon. Bull. Rev. 13, 1–21 (2006).

46. Yonelinas, A. P. The Nature of Recollection and Familiarity: A Review of 30 Years of Research. J. Mem. Lang. 46, 441–517 (2002).

47. Yonelinas, A. P., Aly, M., Wang, W.-C. & Koen, J. D. Recollection and familiarity: examining controversial assumptions and new directions. Hippocampus 20, 1178–1194 (2010).

48. Clark, S. E. & Gronlund, S. D. Global matching models of recognition memory: How the models match the data. Psychon. Bull. Rev. 3, 37–60 (1996).

49. Cleary, A. M. Recognition memory, familiarity, and déjà vu experiences. Curr. Dir. Psychol. Sci. (2008).

50. McNeely-White, K. L. & Cleary, A. M. Music recognition without identification and its relation to déjà entendu: A study using ‘Piano Puzzlers’. New Ideas Psychol. 55, 50–57 (2019).

51. Ford, J. H., Addis, D. R. & Giovanello, K. S. Differential neural activity during search of specific and general autobiographical memories elicited by musical cues. Neuropsychologia 49, 2514–2526 (2011).

52. Ferreri, L., Bigand, E. & Bugaiska, A. The positive effect of music on source memory. Music Sci. 19, 402–411 (2015).

53. Wildschut, T., Sedikides, C., Arndt, J. & Routledge, C. Nostalgia: content, triggers, functions. J. Pers. Soc. Psychol. 91, 975–993 (2006).

54. Schneider, S. Extracting Response Style Bias From Measures of Positive and Negative Affect in Aging Research. J. Gerontol. B Psychol. Sci. Soc. Sci. 73, 64–74 (2017).

55. Jakubowski, K. & Ghosh, A. Music-evoked autobiographical memories in everyday life. Psychology of music (2021).

